# Rebound Excitation of Epileptiform Activities by Transcranial Focused Ultrasound Stimulation

**DOI:** 10.1101/2022.04.30.490021

**Authors:** Taewon Choi, Minseok Koo, Jaesoon Joo, Taekyung Kim, Young-Min Shon, Jinhyoung Park

## Abstract

**Background:** Conventional neurostimulations to treat epilepsy have adverse effects caused by post-inhibitory rebound excitations. Although ultrasound brain stimulation is feasible in inducing anticonvulsant effects, its association with paradoxical rebound excitations is unknown.

**Objective:** This study aimed at demonstrating rebound excitations with transcranial focused ultrasound. The modulations of epileptiform activities toward both suppressive and excitatory responses were investigated by changing ultrasonic transmit sequences.

**Methods:** In a pentylenetetrazol-injected acute epilepsy rat model, transcranial focused ultrasound stimulation was applied on the thalamus to modulate epileptiform activities. The parameters differentiated for pulse sequences were the pulse length, pulse pressure, and interval between the pulses. Sonication effects were assessed by electroencephalography (n=38), immuno-histochemical analysis (n=24), and optical measurement of cerebral blood volume changes (n=18).

**Results:** While ultrasonic patterns of stimuli at long intervals showed antiepileptic effects on electroencephalography, those at short intervals showed rebound excitatory responses followed by inhibitory activities. Further, suppressive states induced by inhibitory stimulations were transformed into excitatory states by applying a consecutive series of short bursts at higher acoustic pressure. Cerebral blood volume changes demonstrated consistent results with electroencephalography. Immunohistochemistry revealed that both inhibitory and excitatory neuronal cells were activated to generate rebound excitatory conditions, while inhibitory cells were activated for suppressive conditions.

**Conclusions:** In our study, variations in ultrasound stimulation patterns could modulate epileptiform activities in both upregulated and downregulated directions.

## Introduction

Epilepsy, a common neurological disorder [1], is caused by failure of an inhibition that disperses epileptic responses over broad brain regions [2]. The main features of antiepileptic therapies, such as antiepileptic drugs (AEDs) [3,4], vagus nerve stimulation (VNS), deep brain stimulation (DBS) [5–7], transcranial magnetic stimulation (TMS), and transcranial direct/alternating current stimulation [5–8], alleviate pathological hyperexcitability by reducing neuronal excitation or enhancing inhibition and are partially effective in suppressing epileptiform activities [9]. Antiepileptic therapies can cause side effects, such as rebound excitations, followed by the application of inhibitory stimuli. Acute rebound effects of AEDs [10,11], VNS, DBS [12–14], and TMS [15,15], are prolonged, probably because of neuronal plasticity [17,18]. The stimulus frequency [13,19,20] and length [21,22] are significant factors inducing rebound. In electrical stimulations, brief stimuli can cause temporary rebound, followed by short inhibition [17,23]. Moreover, fast successive stimulations could exhaust GABA and reinforce excitatory effects, ultimately breaking inhibitory functions [2].

Low-intensity focused ultrasound exerted antiepileptic effects in drug-induced small animal models of temporal lobe epilepsy after stimulating seizure-triggering regions of the anterior thalamic nuclei (ATN) [8] and hippocampus [8,9,21–25]. Despite successful induction of suppressive effects, functional adverse effects are unknown. Because ultrasound stimulation works on inhibitory pathways by activating GABAergic neurons or blocking ionic channels similar to other conventional stimulant methods [24,25,27,28], rebound excitation could be induced by transcranial focused ultrasound (tFUS) stimulation similar to other stimulation approaches.

This study aimed at discovering the ultrasonic stimulus sequences that could upregulate or downregulate epileptiform activities in a pentylenetetrazol (PTZ)-induced acute epilepsy model [29]. Referring to the rebound excitation-triggering electrical stimulations, the ultrasound pulse length, frequency, and intensity were varied to induce paradoxical excitatory responses. Changes in epileptic activities were assessed with electroencephalography (EEG) and optical measurement of cerebral blood volume changes. After each trial, immunohistochemistry was performed to visualize immune responses related to excitatory and inhibitory activations.

## Materials and Methods

All animal experiments conformed to the Korean Ministry of Food and Drug Safety Guide for the Care and Use of Laboratory Animals and were conducted according to the protocols approved by the Institutional Animal Care and Use Committees of Samsung Biomedical Research Institute and Sungkyunkwan University.

### Animal Preparation and Epileptic Induction

The experimental procedure involved a total of 59 male Sprague-Dawley rats, 8 weeks old, weighing 300-320 g (Orient Bio, Inc.). For EEG, eight rats were chosen for the non-stimulated PTZ+/tFUS-control (PTZ-control) group, and five rats were chosen for each stimulated experimental group. Immunohistochemistry was performed within the EEG group for three rats per PTZ-control group and stimulated experimental groups, while three additional rats were evaluated as the PTZ-/tFUS-control (normal control) group. For wide-field optical imaging, three rats in each of the PTZ-control, normal control, and stimulated experimental groups were evaluated. The animals were anesthetized using urethane (1.25 g/kg body weight in 1.25 g/4 mL saline; Sigma-Aldrich, U-2500-500G), and pentylenetetrazol (100 mg/kg body weight in 10 mg/0.1 mL saline; Sigma-Aldrich, P5500-25G) was administered acutely and intraperitoneally to induce epileptic seizures.

### Utrasound Transducer Fabrication

A lead zirconate titanate (DeL Piezo Specialties LLC., DL-47) 690 kHz single-element ring transducer with inner and outer diameters of 10 and 31 mm, respectively, was fabricated with a focal distance of 45 mm and bandwidth of 25%. An acoustic coupling gel-filled cone-shaped beam guide was attached to the frontal side of the transducer. The beam was focused 5 mm ahead of the tip of the collimator with −3 dB beam sizes of 4.7 and 2.2 mm along the axial and lateral directions, respectively (Figs. 1A, 1C, and 1D). By placing the tip of the transducer above the parietal bone (Fig. 1B), the acoustic beam was delivered to the right ATN (A = −1.5 mm, L = +1.5 mm, V = 5.0-5.5 mm; A denotes the anterior (+) or posterior (-) distance from the bregma; L denotes the lateral distance from the bregma; and V denotes the dorsoventral distance from the bregma). The beam pathway was confirmed by overlaying the beam profile on the Paxinos rat brain atlas [30] along the sagittal and coronal planes (Figs. 1E and 1F).

**Fig. 1.**
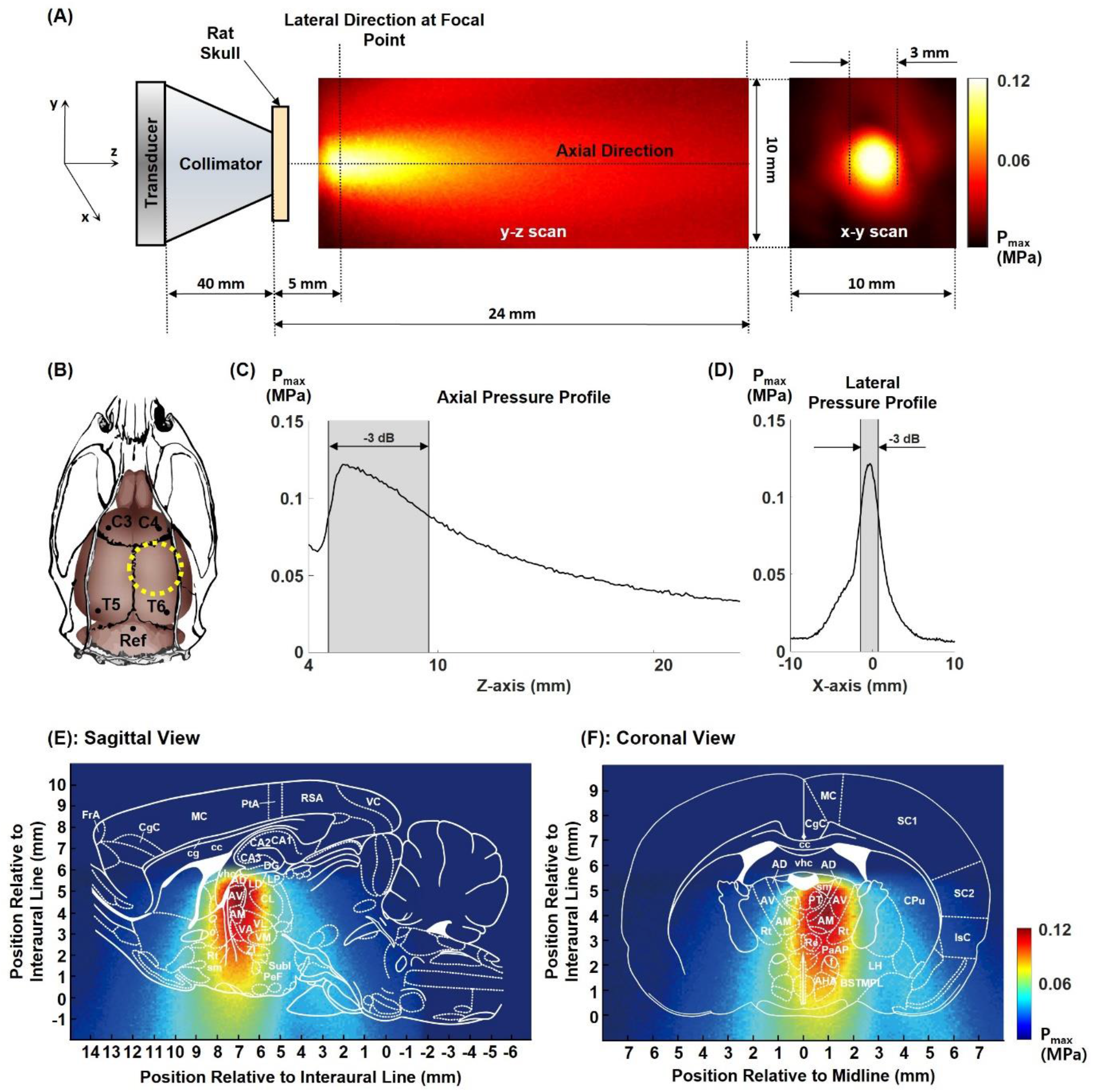
Ultrasound transducer set up for the stimulation. **A** Acoustic beam profile of the ultrasound transducer. **B** Transducer tip and electrode positions in reference to the rat brain and skull. **C, D** Axial and lateral pressure profile of the acoustic beam. **E, F** Sagittal and coronal views of the ultrasound beam superimposed to the Paxinos rat brain atlas [30].

### Sonication and Monitoring Protocol

To simplify the ultrasonic conditions and mimic electrical stimulations generating rebound excitations, the stimuli were classified into two types: one with low pressure (0.25 MPa) consecutive burst train (TX1-TX4 in Fig. 2A) and the other with high pressure (1.0 MPa) with shorter bursts (TX5 in Fig. 2A) with a 50% duty cycle of 100 Hz pulse repetition frequency. TX6, a back-to-back transmission of TX3 and TX5, was generated with the assumption that rebound excitation could be artificially induced using brief excitation followed by inhibitory stimulation.

**Fig. 2.**
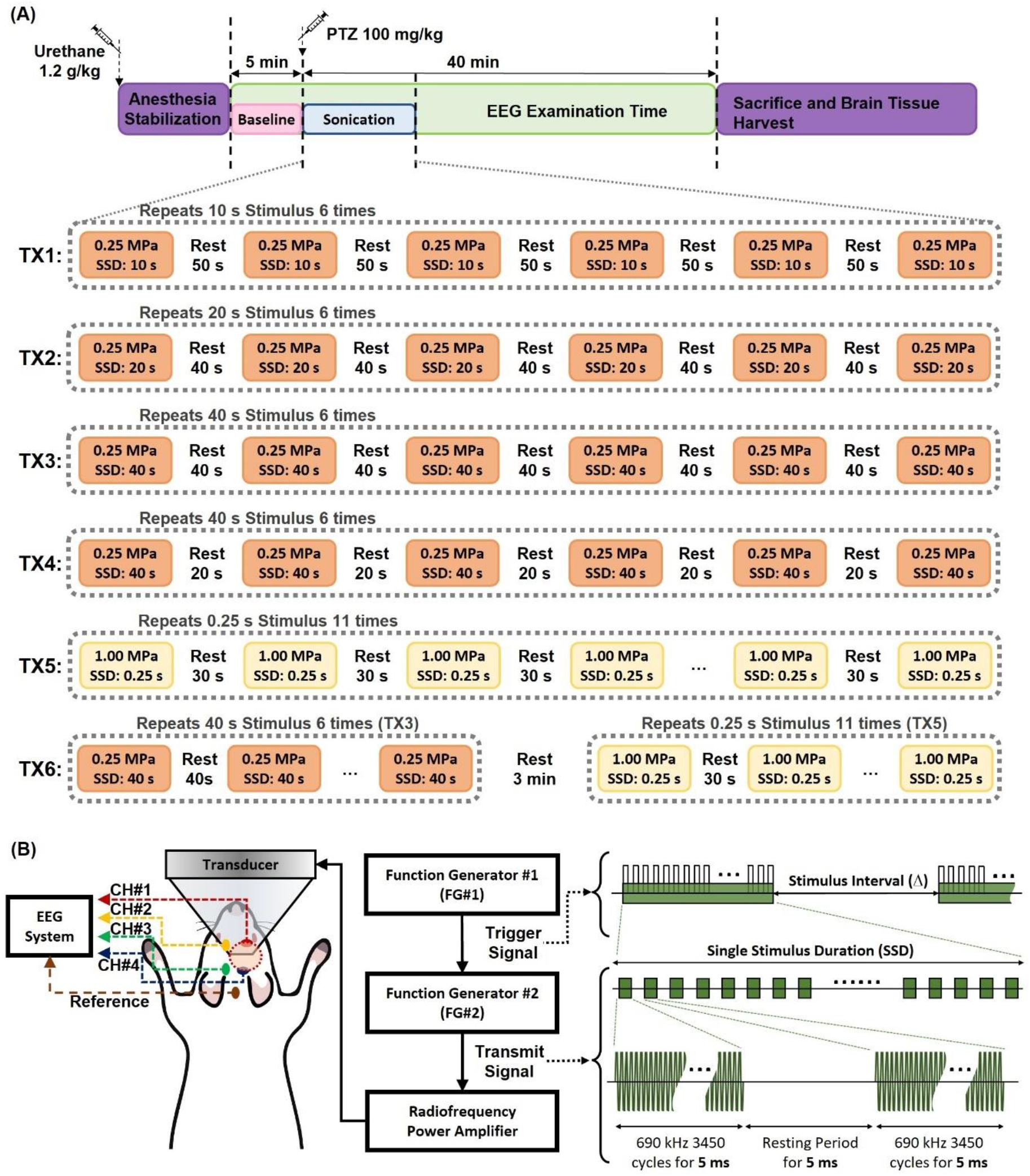
Focused ultrasound pulse sequences and pulse platform for stimulation. **A** Sonication protocols of TX1-TX6 stimulations. The modulation effects were explored by changing the duration of each low-pressure burst to 10 (TX1), 20 (TX2), and 40 (TX3 and TX4) s. The number of bursts was six for each stimulation condition, and the burst intervals for sequences from TX1 to TX4 were 50, 40, 40, and 20 s, respectively. A high-pressure burst train (TX5) was transmitted for 5 min during which the length of the burst was 250 ms with a transmit interval of 30 s. The pulse sequence of TX6 represents the back-to-back transmissions of TX3 and TX5 with 3-min resting between the two sequences. **B** Overall experimental system for ultrasound stimulations with simultaneous four-channel EEG recording and signal shape generated from each function generator to obtain the stimulation pulse sequence.

Fig. 2B depicts the overall experimental platform to deliver ultrasonic energy to the target region and record EEG signals. While the rats were anchored to a custom-made stereotaxic frame (Scitech Korea, Inc.), ultrasound pulsation was generated using two function generators. The first function generator (Tektronix, AFG3252) sent out a pulse train of 100 Hz, and the connected second function generator (Agilent, 33250A) transmitted a 5-ms, 3,450-cycle, 690-kHz burst to secure a 50% duty cycle (Fig. 2B). The generated waveform was then passed through a radiofrequency power amplifier (Electronics and Innovation, 350 L) and applied to the transducer. The experimental protocol involved: 1) anesthesia; 2) pre-tFUS (baseline) monitoring; 3) PTZ injection and sonication; 4) post-tFUS monitoring; and 5) sacrifice and brain tissue harvest. Supplementary Fig. 2 shows the overall timeline for sonication and data acquisition.

### EEG Recording

To record EEG signals, rats were stereotactically implanted with five cortical screw electrodes (Fine Science Tools, 19010-00 screws). The implantation coordinates were adopted from a stereotactic atlas [30,31]: C4/C3 (primary motor cortex; A = +2.2 mm, L = ±3.2 mm) and T6/T5 (temporal association cortex; A = −8.3 mm, L = ±5.8 mm). The reference ground electrode was implanted on the posterior side of the lambda. EEG was performed to monitor neuronal activity changes using a four-channel commercial EEG acquisition system (Natus, NicoletOne vEEG) at a sampling rate of 500 Hz. To assess the epileptic severity, the ratio of the number of epileptic spikes at baseline with the ones at specific timepoints was calculated. Epileptic spikes were defined as the local maxima points with amplitudes larger than twice the standard deviation from individual baseline EEG activities lasting 70 ms.

### Immunohistochemistry

Immunohistochemistry was performed by conducting c-Fos, glutamic acid decarboxylase 65-kilodalton isoform (GAD65), ionized calcium-binding adaptor molecule 1 (Iba-1), and glial fibrillary acidic protein (GFAP) staining of rat brain sections. The rats were sacrificed after EEG recording and transcardially perfused with saline to obtain brain tissue samples. To compare the degree of excitatory responses generated under ultrasonic stimulus conditions (TX1 TX6), the ratio of c-Fos-positive cells to total cells in the granular layer of the dentate gyrus was quantified using ImageJ software (National Institutes of Health, 1.53c). To compare inhibitory synaptic transmission, GAD55-positive cell densities were measured in the molecular and granular layers of the dentate gyrus. Iba1 and GFAP analyses were performed to investigate the reactivity of microglia and astrocytes, respectively, by computing the Iba1- and GFAP-positive cell densities.

### Cerebral Blood Volume Measurement

Craniotomy was performed on isoflurane-anesthetized rats secured in a custom-made stereotaxic frame. The cranial window region was generated above the left ATN (A = −1.5 mm, L = −1.5 mm) such that the tFUS stimulus could be applied on the right ATN while simultaneously imaging the cranial window. After an overnight recovery, urethane (1.25 mg/kg body weight) was injected intraperitoneally for wide-field optical imaging.

To examine the regional hemodynamic response of cerebral blood volume (CBV), optical measurement was conducted for the normal control, PTZ-control, and tFUS-stimulated groups. The imaging system comprised a macro-zoom microscope (Olympus, MVX10), an sCMOS camera (Andor, Zyla 5.5), and a light-emitting diode source. For the experiment, raw reflectance data were collected under green (530 nm) and red (625 nm) light, as they are sensitive primarily to changes in the local total blood volume, denoted as [HbT], and the local concentration of deoxyhemoglobin, denoted as [HbR], respectively [32–34]. Each imaging trial was recorded for 30 min, including a 2-min baseline (pre-PTZ/tFUS).

### Statistical Analyses

All experimental data were statistically analyzed using IBM SPSS Statistics, Version 27 (Armonk, NY: IBM Corp). Tests used for the statistical analyses are presented along with the *p*-values throughout the paper. *P*-values less than 0.05 were considered statistically significant.

## Results

Fig. 3 shows the representative EEG signals for all transmit sequences. While the sparsely located peaks were identified at baseline, the signal was transformed into ictal shapes that showed repetitive high frequencies discharging peaks in the PTZ-control group. With TX1 stimulation, the peak oscillation frequency was higher compared to the PTZ-control group. Discharging activities in TX3 became almost similar to the baseline curve, which intermittently presented discharging spikes. With TX4 stimulation, the form of discharging activities resembled that at baseline but showed more frequent and dense spikes with higher amplitudes overall. With TX5 stimulation, the discharging pattern gradually became comparable with that of the PTZ-control group.

**Fig. 3.**
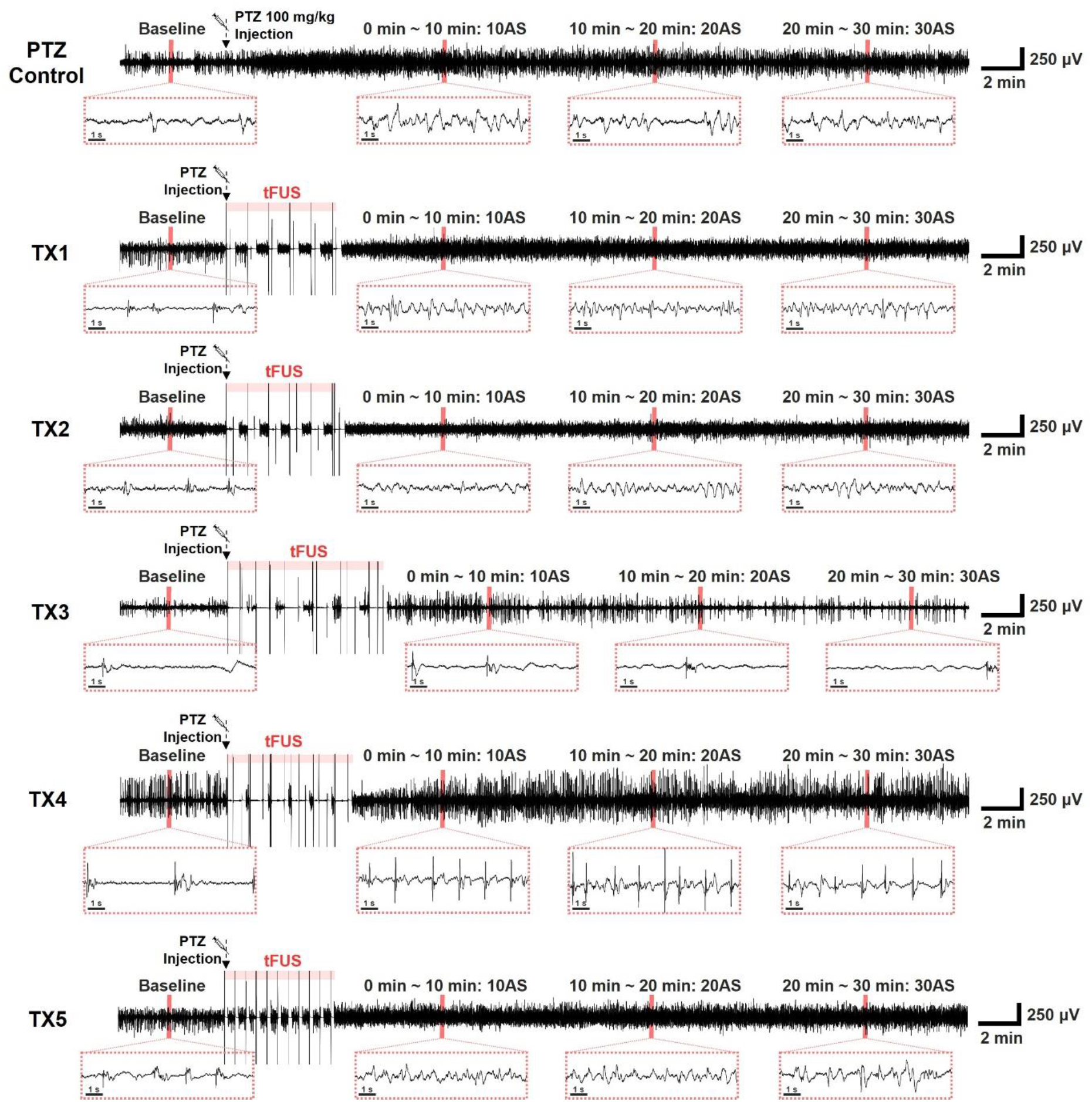
Six representative EEG waveforms. The signals from the first to sixth rows present the time courses of EEG signals in the PTZ-control and stimulated experimental groups using the transmit conditions from TX1 to TX5, respectively. The graphs under the time courses are the magnified representative signals during the resting baseline period and 0–10 (10AS), 10–20 (20AS), and 20–30 (30AS) min after the stimulation.

Fig. 4A shows the EEG time course plot with the TX6 stimulus. The baseline plot resembles the shape of the curve followed by TX3, which barely showed discharging peaks or repetitive patterns. In contrast, the silent signal changed to repetitive spikes, which could be identified in the magnified plots at 10AS and 20AS. Fig. 4B shows the number of epileptic spikes from the start of stimulations. With TX3 alone, the number of epileptic spikes did not change significantly from baseline even after drug administration. However, the number of spikes sharply increased from the end of TX5 by almost thrice higher than that of the PTZ-control group. There was a rebound overshoot at 15-20 min, and the rebound excitatory effect lasted further until 30-35 min (*p* = 0.913; two-tailed paired *t*-test) and maintained a slightly higher number of discharging peaks compared to the PTZ-control group (*p* = 0.087 at 30-35 min; one-tailed Mann-Whitney U test).

**Fig. 4.**
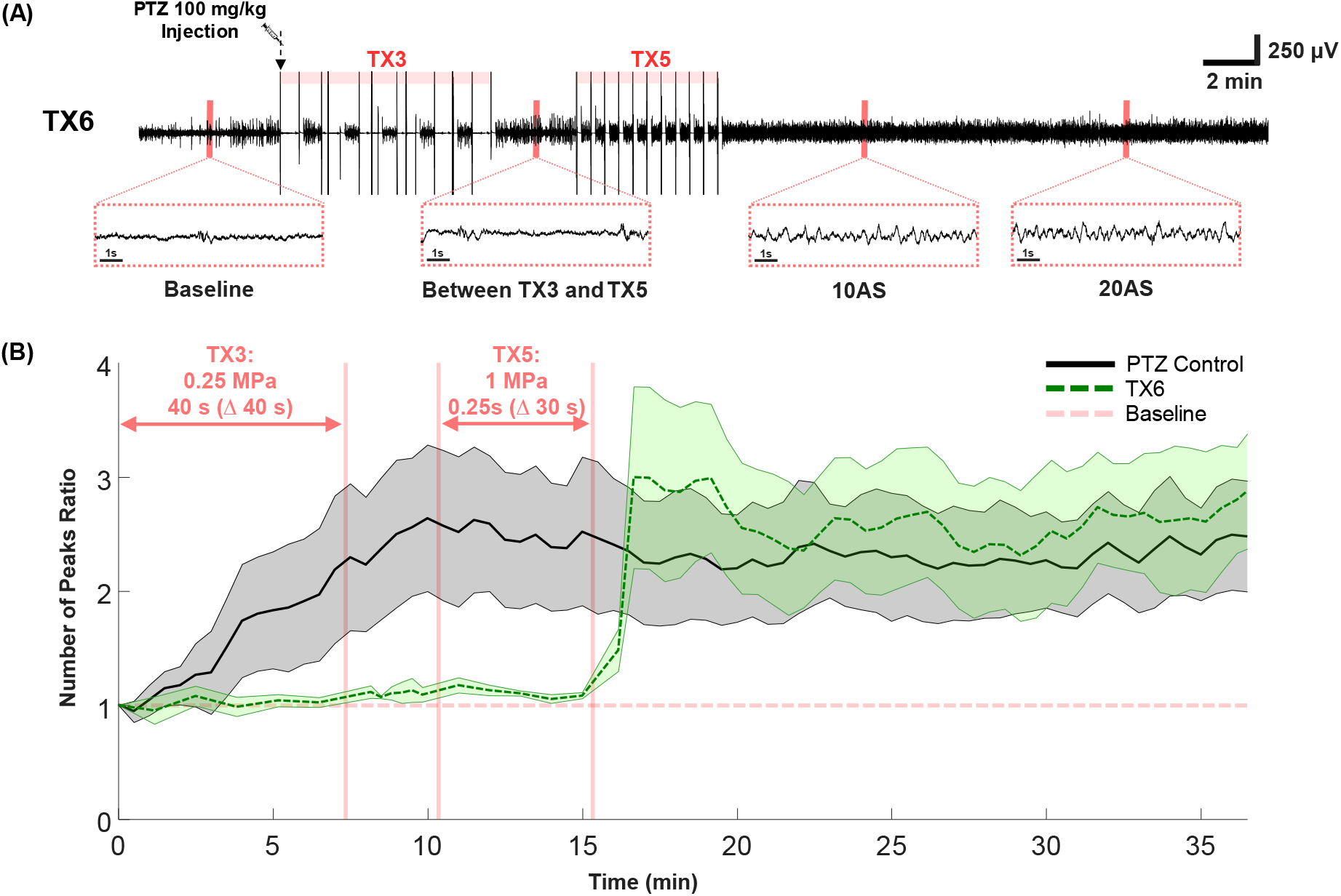
EEG study results from the group stimulated with TX6 (consecutive transmits of TX3 and TX5). **A** Representative EEG discharging curve acquired for 35 min and magnified plots at baseline, the resting time between TX3 and TX5, 10AS and 20AS. **B** The number of epileptic spikes from the start of TX3 to the end of recording. The black solid line and green dashed line show the ratios in the PTZ-control and TX6 groups, respectively.

Fig. 5A shows the comparison of the severity of epilepsy based on the number of discharging peaks in EEG signals from the C4 electrode. Over a post-stimulation period of 20 min, the number of epileptic spikes in the PTZ-control group increased by more than twice compared to baseline. Although the level slightly decreased from 5AS to 20AS in the PTZ-control group, the change was not statistically significant (*p* = 0.764; two-tailed paired *t*-test). As the stimulation length elongated from 10 to 40 s (from TX1 to TX3), the number of epileptic spikes decreased (*p* = 0.003, 0.005, and 0.624 at 20AS for TX1, TX2, and TX3 compared to baseline, respectively; two-tailed paired *t*-test) and ultimately became comparable to baseline with TX3 (*p* = 0.515, 0.995, 0.478, and 0.624 at 5AS, 10AS, 15AS, and 20AS, respectively; two-tailed paired *t*-test). In contrast, TX4 with shortened burst intervals generated a comparable number of epileptic spikes as the PTZ-control group (*p* = 0.457, 0.185, 0.232, and 0.061 at 5AS, 10AS, 15AS, and 20AS, respectively; one-tailed Mann-Whitney U test). Moreover, the increase in the number of epileptic spikes with TX1 was statistically significant compared to that in the PTZ-control group at the post-stimulation periods from 5AS to 20AS (*p* = 0.083, 0.031, 0.061, and 0.014 at 5AS, 10AS, 15AS, and 20AS, respectively; one-tailed Mann-Whitney U test). With TX6 consecutively transmitting TX3 and TX5, the number of peaks increased by 12.4% more than that with TX1 alone at 5AS (*p* = 0.0014; one-tailed Mann-Whitney U test), but the level afterward became similar to that with TX1 (*p* = 0.009, 0.002, and 0.403 at 10AS, 15AS, and 20AS, respectively; one-tailed Mann-Whitney U test). In addition, the number of epileptic spikes with TX6 was always higher compared to the PTZ-control group over all post-stimulation periods at 10AS, 15AS, and 20AS, although the differences were not statistically significant *(p* = 0.145, 0.285, and 0.083, respectively; one-tailed Mann-Whitney U test).

**Fig. 5.**
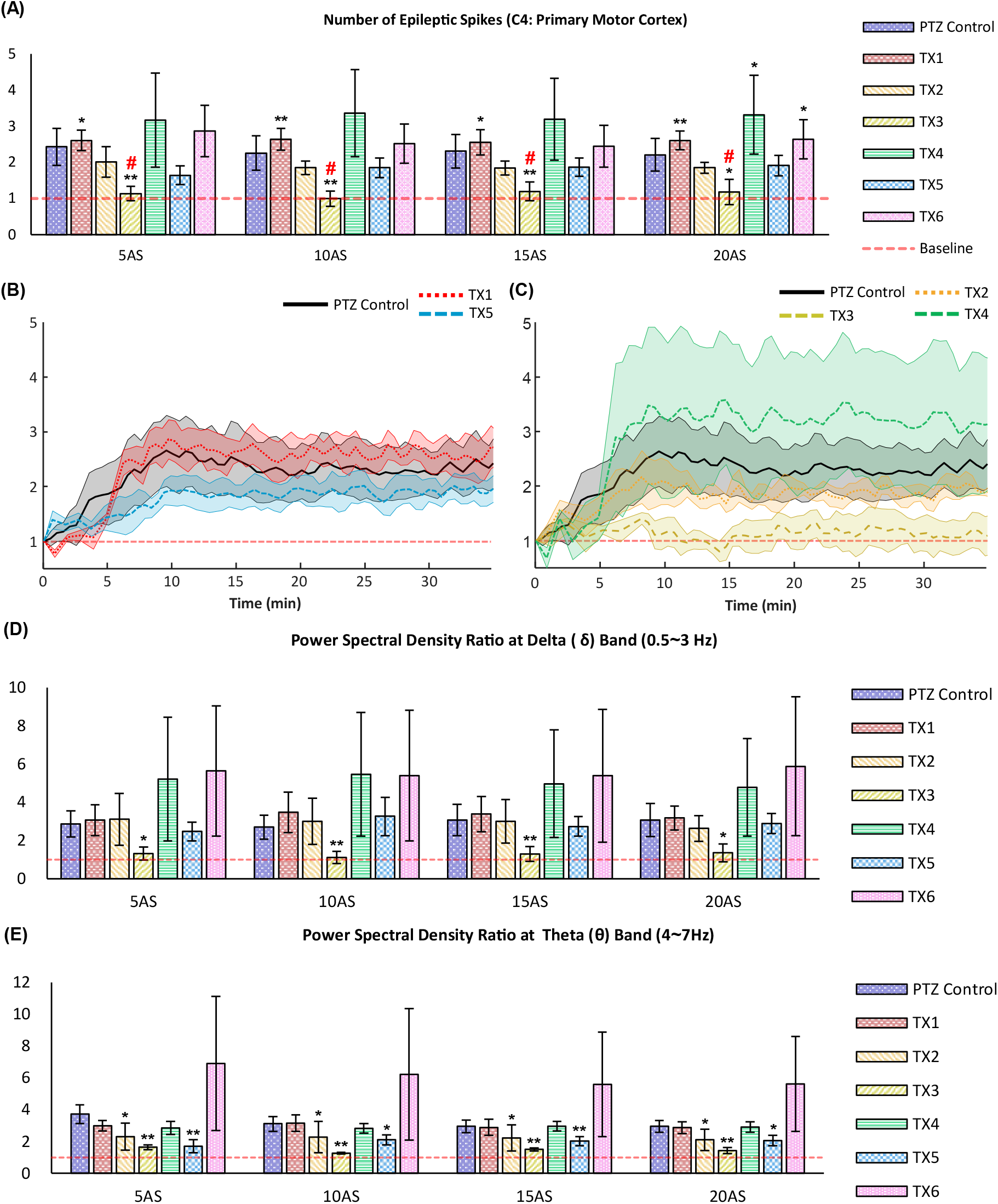
Assessment of the severity of epileptic seizures based on the number of epileptic spikes and power spectral density measurements. The ratio of the number of epileptic spikes in each transmit condition was measured at time periods of 5 (5AS), 10 (10AS), 15 (15AS), and 20 (20AS) min after stimulation. The red-dashed transverse line in the middle of each graph indicates the baseline level. \**p* < 0.10 against the PTZ-control group (one-tailed Mann-Whitney U test). \*\**p* < 0.05 against the PTZ-control group (one-tailed Mann-Whitney U test). ^#^*p* > 0.10 against baseline (two-tailed Mann-Whitney U test). **A** Number of epileptic spikes measured from the C4-Ref EEG signal. **B, C** Time course of the ratio in each group. **D, E** Power spectral density ratio of EEG signals at the delta (0.5-3 Hz) and theta (4-7 Hz) frequency bands.

**Fig. 6.**
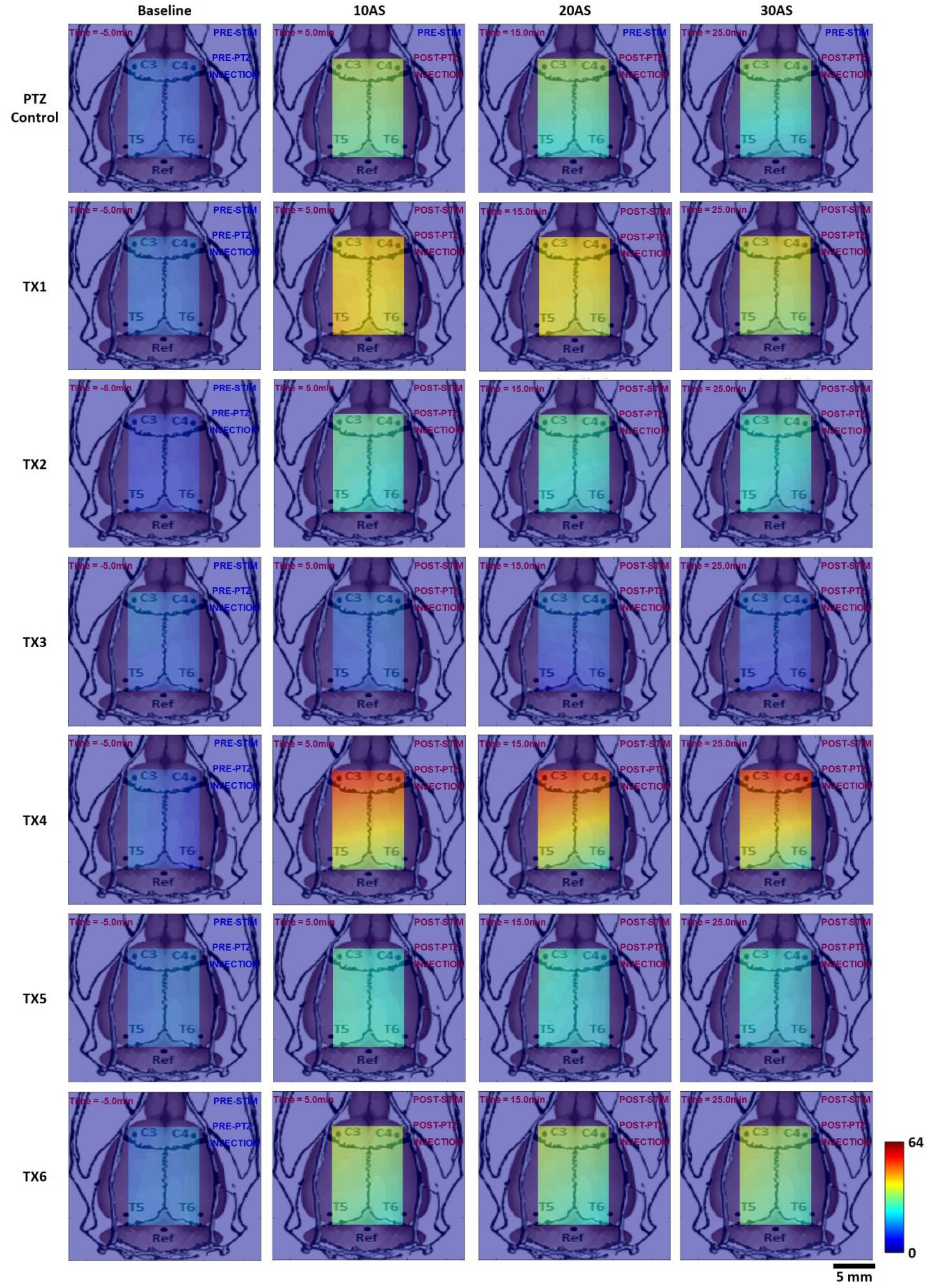
Two-dimensional topographical maps representing the number of epileptic spikes by time measured at electrodes in C3, C4, T5, and T6. The color map images were acquired by interpolating the number of epileptic spikes acquired with EEG signals from the four electrodes. The measured numbers were placed at the four corners of the topological window, and the twodimensional information was calculated by two-dimensional linear interpolation. The colorized scale is shown in the color map at the bottom right of the figure. The representative images at baseline and 10 (10AS), 20 (20AS), and 30 (30AS) min after stimulus are presented. Supplementary Vids. 1 - 7 are the movies in the PTZ-control and TX1-TX6 groups, respectively.

Figs. 5B and 5C show the time courses of the number of epileptic spikes in the post-drug injection periods. TX1 exhibiting rebound excitatory effects in the post-stimulus period showed suppressive effects at a lower level compared to the PTZ-control group in the stimulus period from 0 to 5 min (Fig. 5B). TX4 stimulation showed rebound excitatory effects in the post-stimulus period but suppressive effects in the stimulus period from 0 to 5 min (Fig. 5C). With TX3 stimulation, the prominent suppressive effects lasted during and after the stimulation.

Figs. 5D and 5E compare the power spectral density (PSD) among control and transmit conditions from TX1 to TX6 in the delta and theta frequency bands, respectively. In the delta band, the spectral power of EEG for the TX1 condition increased higher compared to the PTZ-control group (*p* = 0.153 at 10AS; one-tailed Mann-Whitney U test), while for the TX3 condition was lower compared to the PTZ-control group (*p* = 0.014 at 10AS; one-tailed Mann-Whitney U test). The elevation of the power spectral level with TX5 was noticeable because the mean was 1.914 times higher than that of the control group and 5.884 times higher than the baseline level at 20AS. In the theta band, the PSD level with TX1 was similar to that of the control after 10AS, while the highest level was presented with TX5 (Fig. 5E). With TX3 stimulation, the level was suppressed to 1.647 as soon as the stimulation was completed and was comparable with the baseline level at 10AS (*p* = 0.001 against PTZ-control, one-tailed Mann-Whitney U test). The suppressive effects lasted until the end of the experiment.

Fig. 5 shows a picture of a rat brain superimposed on the two-dimensional topographical map images that visualize the degree of brain activities throughout the brain area by time. The combined image was generated every 30 s until the end of acquisition, and time-series images were converted into a movie for each transmit condition. Supplementary Vids. 1-7 are for the PTZ-control and TX1-TX5 groups, respectively. While the patterns of amplitude change were comparable to the results shown in Figs. 4 and 5, the number of peaks was higher in the frontal cortices in C3 and C4 than in the central cortices in T5 and T5. The patterns in both hemispheres were similar.

Fig. 7A shows c-Fos immunohistochemistry results of the samples harvested in the sacrificed animals immediately after each experiment. While the expressed cells were rarely found in the normal control group, c-Fos-labeled cells were abundant in the PTZ-control. Proportions (22.5%) of c-Fos-positive cells were higher with TX1 compared to the PTZ-control group (n = 3; *p* = 0.100; one-tailed Mann-Whitney U test). In contrast, only a couple of expressed cells in the histology could be discovered with TX3. As shown in the bar graph, 80.72% (standard error of mean [SEM] = 2.00%) of neuronal cells were c-Fos expressed with TX1, while 71.73% (SEM = 7.12%) of cells were presented in the PTZ-control group. TX2 (*p* = 0.303; one-tailed Mann-Whitney U test) and TX4 (*p* = 0.855; one-tailed Mann-Whitney U test) showed c-Fos-positive cell ratios comparable to those in the PTZ-control group. Additionally, TX5 showed a comparable ratio to TX1 (mean = 80.05%; SEM = 0.63%), and TX6 presented an even higher value of approximately 87.27% (SEM = 0.88%) than TX1 (*p* = 0.021; one-tailed Mann-Whitney U test). However, the expression ratio dramatically reduced to 24.22% with TX3.

**Fig. 7.**
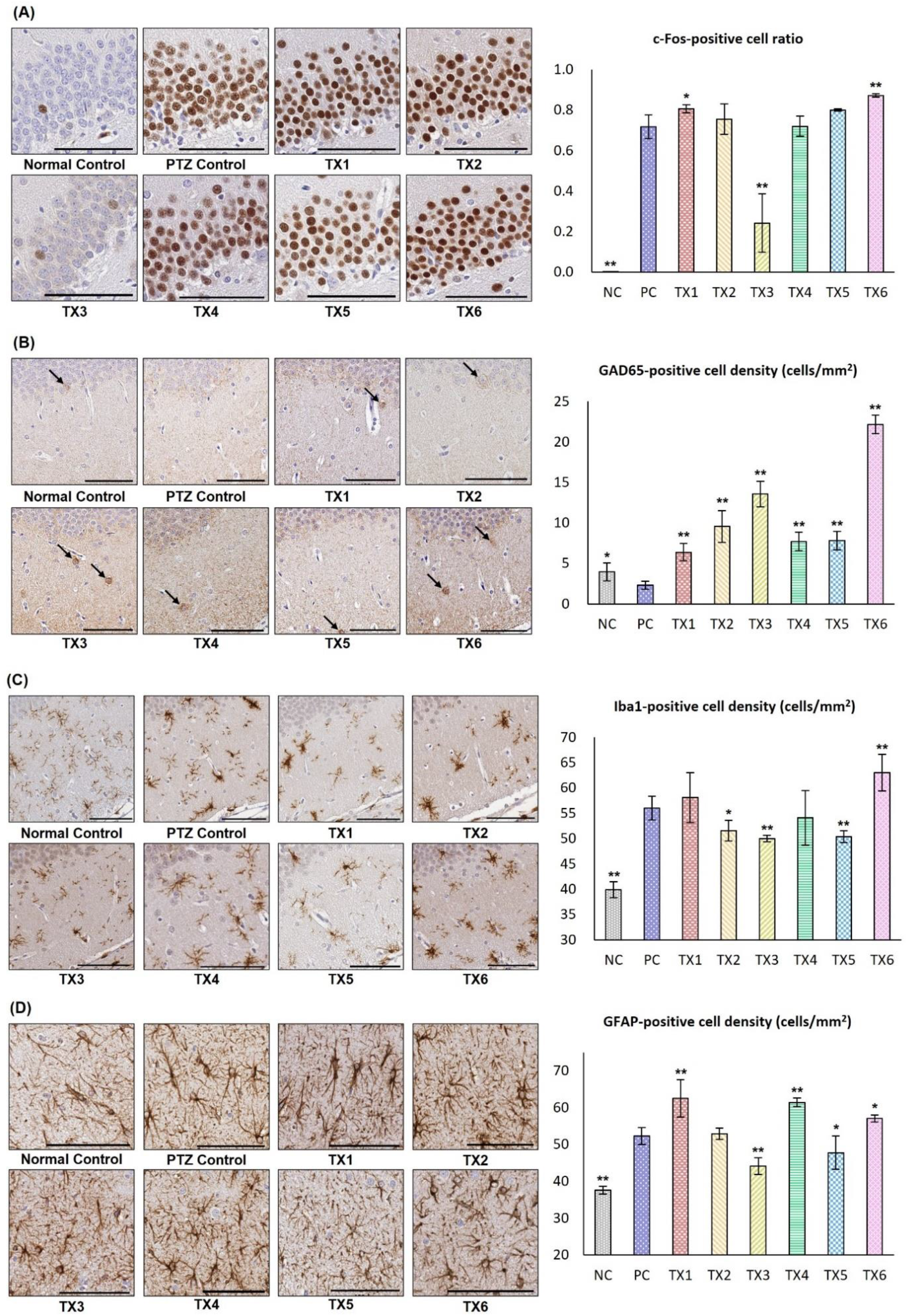
Representative brain slice images and quantitative measurements from immunohistochemistry. “NC” denotes normal control, and “PC” denotes PTZ -control. The scale bar denotes 100 μm for all representative images. *\*p* < 0.10 against the PTZ-control group (one-tailed Mann-Whitney U test). \*\**p* < 0.05 against the PTZ-control group (one-tailed Mann-Whitney U test). A Representative images of the c-Fos-stained brain slice at the dentate gyrus and quantitative measurements of the stained cell ratio for c-Fos. B Representative images of the GAD65-stained brain slice at the dentate gyrus and quantitative measurements of GAD65-positive cell density. C Representative images of the Iba1-stained brain slice at the dentate gyrus and quantitative measurements of Iba1-positive cell density. D Representative images of the GFAP-stained brain slice at the dentate gyrus and quantitative measurements of GFAP-positive cell density.

Fig. 7B shows GAD65 immunohistochemistry results. While both normal and PTZ-control cases barely showed GAD65-positive cells, the other conditions presented more expressed cells. The GAD65-positive cell density with TX3 (mean = 13.58 cells/mm^2^; SEM = 3.16 cells/mm^2^) was approximately 5.8 times higher than that in the PTZ-control group (mean = 2.33 cells/mm^2^; SEM = 0.98 cells/mm^2^). Although sharing the same duration for the single stimulus, TX4 showed lower expression of GAD65-positive cells than TX3 (*p* = 0.017; one-tailed Mann-Whitney U test) but higher expression compared to the PTZ-control group (*p* = 0.017; one-tailed Mann-Whitney U test). The GAD65-expressed cell density with TX1 (mean = 6.41 cells/mm^2^; SEM = 1.84 cells/mm^2^) was less than the density with TX4 (mean = 7.82 cells/mm^2^; SEM = 1.97 cells/mm^2^), while both were lower than the one with TX3. The GAD65-expressed density with TX6 (mean = 22.16 cells/mm^2^; SEM = 14.14 cells/mm^2^) was the highest, which seems to be close to the summation of the densities with TX3 and TX5 (sum of means = 21.40 cells/mm^2^).

Fig. 7C shows the immunoreactivity of Iba1, with the expressed cells according to a previous study [35]. While the Iba1-expressed cell density in the normal control group was 39.93 cells/mm^2^, the number increased to 56.03 cells/mm^2^ in the PTZ-control group and was to 58.11 cells/mm^2^ with TX1. The activated cell density with TX3 was 50.02 cells/mm^2^, only higher than the normal control group. TX4 increased the expression of Iba1 to 54.13 cells/mm^2^, higher than that with TX3. With TX5, the expression level was 50.39 cells/mm^2^, which was comparable to that with TX3 (*p* = 0.533; two-tailed Mann-Whitney U test). TX6 induced the highest Iba1-expression level at 63.04 cells/mm^2^ among the experimental groups.

Fig. 7D shows the immunoreactivity of GFAP indicating astrocytic reactivity. GFAP density increased to 52.35 cells/mm^2^ in the PTZ-control group, unlike the normal control group, which presented a density of 37.61 cells/mm^2^. The GFAP-positive cell densities of TX1, TX2, and TX3 gradually decreased to 52.50, 52.95, and 44.15 cells/mm^2^, respectively. TX4 showed a higher density of 61.45 cells/mm^2^ compared to TX3 (*p* = 0.025; one-tailed Mann-Whitney U test). The degree of GFAP expression with TX5 was similar to that with TX3, showing a reduced level of 47.78 cells/mm^2^ (*p* = 0.413; one-tailed Mann-Whitney U test). TX5 presented a comparable GFAP-density level of 57.06 cells/mm^2^ with TX1 and TX4, which was as high as the density in the PTZ-control group (*p* = 0.050; one-tailed Mann-Whitney U test).

Fig. 8A shows the cranial window for the optical measurement, and Fig. 8B shows the changes in CBV, which are coupled with neuronal activities [37]. While Δ[HbO] barely changed in the normal control group, it sharply increased in the PTZ-control group after the drug administration. With TX1, Δ[HbO] sharply increased +19% compared to baseline and was maintained at a higher level compared to the Δ[HbO] in the PTZ-control group. The increase in Δ[HbO] was suppressed using TX3, and the Δ[HbO] level was maintained at a level lower than that in the PTZ-control group. With TX5, Δ[HbO] showed a brief overshoot in the stimulation period, followed by a gradual decay. With TX6 and consecutive transmissions of TX3 and TX5, increase in the Δ[HbO] level in the PTZ-control group was minimized to approximately +7.5% from baseline, and the suppressive response was translated to the excitatory state after completion of sonication with TX5, further increasing to approximately +16% from baseline at 30 min.

**Fig. 8.**
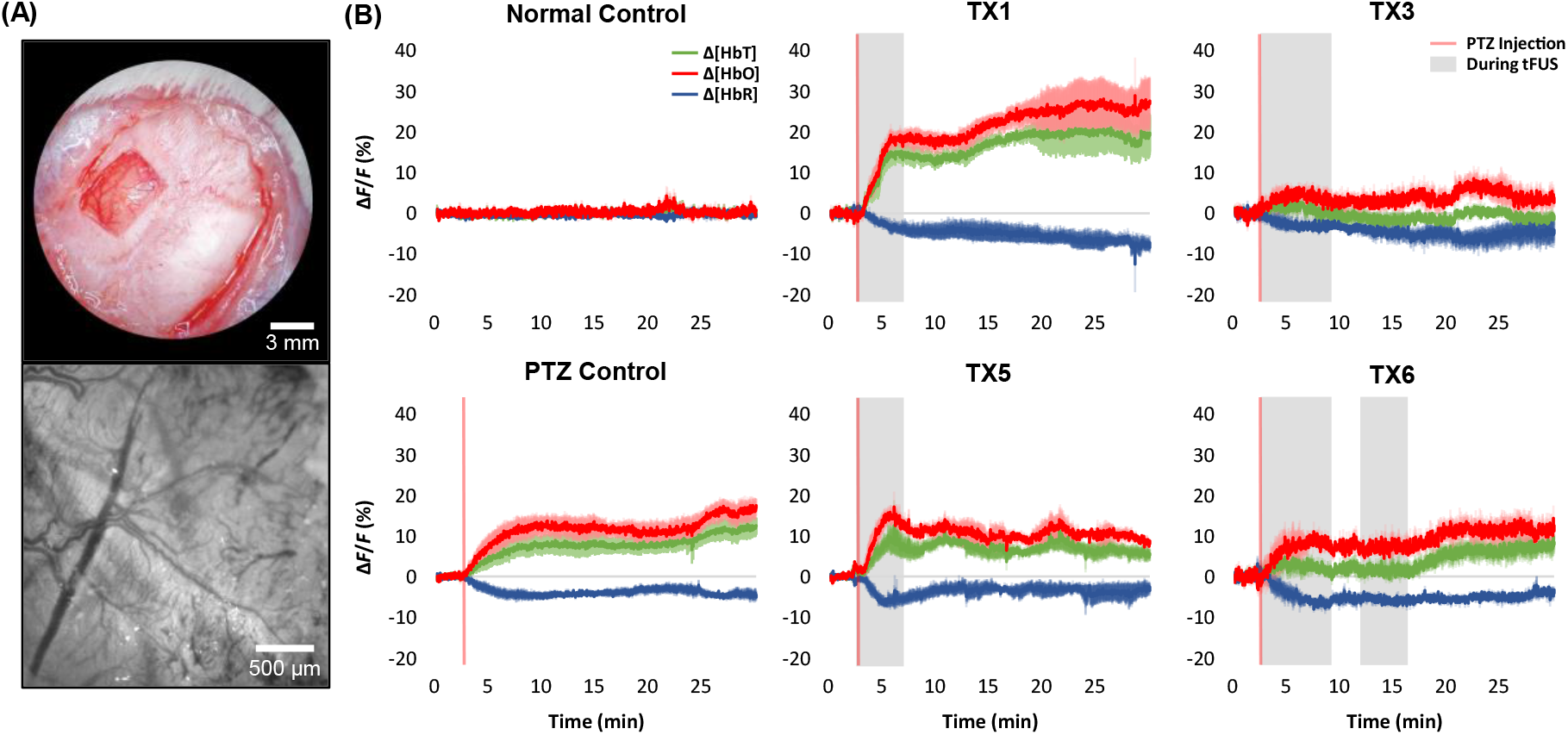
Cerebral blood volume measurement on wide-field optical imaging. “[HbT]” denotes the total hemoglobin concentration (in green); “[HbO]” denotes the oxyhemoglobin concentration (in red); and “[HbR]” denotes the deoxyhemoglobin concentration (in blue). The vertical red line denotes the time of PTZ injection and tFUS initiation. **A** Photographs of the fabricated cranial window and visualized cortical surface of the rat brain acquired by a stereoscopic microscope and a macro zoom microscope at top and bottom, respectively. **B** Changes in cerebral blood volume calculated from reflectance data.

## Discussion

Despite successful demonstrations of lowering the epileptiform activities with tFUS in previous studies, possible adverse effects of worsening the disease after suppressive stimulations were not considered. As previous studies using electrical stimulations have reported rebound excitation [13,19–23], variation in the length of the ultrasound burst could modulate epileptiform activities between suppressive and excitatory states. Rebound excitation was identified after ultrasound stimulation of 0.25 MPa for 10 s based on EEG and CBV results. Because rebound excitation includes both inhibitory and excitatory responses in series, inhibitory responses might have caused the increase in GAD65 during TX1 stimulation, and the elevated c-Fos could result from rebound excitation. Rebound excitation with TX1 seemed to induce more brain damage compared to the PTZ-control group as more Iba1- and GFAP-positive cells were identified with TX1 stimulation in the hippocampal region compared to the PTZ-control group (Fig. 7).

In contrast, as the burst length increased to 40 s with a burst interval of 40 s, only suppressive effects could be identified for at least 30 min after stimuli. Unlike TX1, TX3 maintained the baseline level during stimulation (Fig. 5C), with no rebound excitation after stimuli until the end of the experiment. The short bursting intervals of 20 s could trigger rebound excitations with the same stimulation length as shown by EEG, CBV, and immunohistochemistry with TX4. Because the excretion time for GABA with electrical stimulation was more than 40 s in a mouse model [38], the interval between ultrasound stimulations might need to be longer than 40 s to maintain the decent GABA level in the synaptic spaces. The reduced stimuli interval would reduce GABA excretion, and excitatory responses could dominate inhibitory responses. In the immunohistochemistry studies, on TX3, the number of c-Fos-expressed cells reduced by 55%, while GAD65 increased by 583% compared to the PTZ-control group (Fig. 7). However, with TX4, the expression of GAD55 was significantly reduced by 43.24% compared to that with TX3 (*p* = 0.017; one-tailed Mann-Whitney U test), while the c-Fos expression was comparable to that in the PTZ-control group.

In addition to inducing rebound effects with TX1 alone, rebound excitation could be generated by combining the suppressive transmission of weak long stimulations with strong short stimulations. The application of TX5, a back-to-back transmission of TX3 and TX5 with a 3-min interval, could induce a transition from suppressive responses to ironically amplifying the epileptiform activities, although TX3 or TX5 alone induced suppressive effects (Figs. 5B and 8B). The elevated low-frequency PSD-level with TX6 might mean that TX6 rendered the EEG signal in the delta and theta bands more synchronized compared to the other transmission sequences during rebound excitation. Considering that the increase in delta and theta band PSD-levels was a signal marker for mesial temporal lobe epilepsy in previous studies [35], epileptiform activities might have worsened more compared to the PTZ-control group. In immunohistochemistry studies, the expression rate of GAD65 was the highest with TX6, in addition to c-Fos. Because immunohistochemistry visualizes the accumulated activated responses at the end of each experiment, combined TX3 and TX5 could generate the expressions of GAD65 and c-Fos. Amplified rebound excitation also increased brain damage more compared to the other conditions. The number of Iba1 expressed cells was the highest with TX6, although the GFAP expression level with TX6 was similar to that in the PTZ-control group.

The number of Iba1-positive cells, which are reactive near-damaged neuronal cells, was sharply increased in the PTZ-control group compared to the normal control group. The amount of activation increased with TX1 and TX6 and decreased with TX3 and TX5. However, the level was higher than that of the normal control group. Therefore, tFUS sequences inducing higher EEG discharges could induce more cellular damage, while the other sequences suppressing epileptiform activities could save neuronal cells although the suppressive treatments could not fully recover the damage, unlike the normal control group. To confirm that the damage in neuronal cells was caused by the proconvulsive effect of tFUS stimulation and not by the delivery of the ultrasonic energy, Cresyl Violet staining was performed on both normal control and PTZ-/tFUS+ groups. The Nissl-stained brain slice images demonstrated no damage in the rat brain after TX3 and TX5 stimulations, which could be the worst cases with respect to sonication time and amplitude, respectively (Supplementary Fig. 1).

## Conclusion

In our study, modulation of epileptiform brain activity was demonstrated by purely changing the pulse duration, interval, and amplitude. The rebound effect worsening epileptiform activities could be induced by applying brief stimulations or decreasing the stimuli interval, which might make the amount of glutamate higher than GABA. In addition, the rebound effect could be amplified by using back-to-back transmissions of the low-pressure-elongated and high-pressure-shortened stimuli, which resulted in worsening of the epileptiform activities followed by fully suppressive effects.

## Supporting information

2D topographical map representing the number of epileptic spikes ratio by time in PTZ-control

2D topographical map representing the number of epileptic spikes ratio by time in TX1

2D topographical map representing the number of epileptic spikes ratio by time in TX2

2D topographical map representing the number of epileptic spikes ratio by time in TX3

2D topographical map representing the number of epileptic spikes ratio by time in TX4

2D topographical map representing the number of epileptic spikes ratio by time in TX5

2D topographical map representing the number of epileptic spikes ratio by time in TX6

## Abbreviations

AD: anterodorsal thalamic nucleus
AED: antiepileptic drug
AHA: anterior hypothalamic area, anterior part
AM: anteromedial thalamic nucleus
AP: anteriorly (+) or posteriorly (-) from the bregma
AS: minutes after stimulations
ATN: anterior thalamic nuclei
AV: anteroventral thalamic nucleus
BSTMPL: bed nucleus of stria terminalis, medial division, posterolateral
CA: cornu ammonis
cc: corpus callosum
cg: cingulum
CgC: cingulate cortex
CL: centrolateral thalamic nucleus
Cpu: caudate putamen (striatum)
DBS: deep brain stimulation
DG: dentate gyrus
DV: dorsoventral distance from the horizontal plane passing through the bregma and lambda on the surface of the skull
EEG: electroencephalography
f: fornix
FG: function generator
FrA: frontal association cortex
GAD65: glutamic acid decarboxylase 65-kilodalton isoform
GFAP: glial fibrillary acidic protein
Iba1: ionized calcium binding adaptor molecule 1
ILAR: Institute of Laboratory Animal Resources
IsC: insular cortex
LD: laterodorsal thalamic nucleus
LH: lateral hypothalamic area
LP: lateral posterior thalamic nucleus
MC: motor cortex
ML: laterally from the bregma
NC: normal control
PaAP: paraventricular hypothalamic nucleus, anterior parvicellular part
PC: pentylenetetrazol-control
PeF: perifornical nucleus
PSD: power spectral density
PT: paratenial thalamic nucleus
PtA: parietal association cortex
PTZ: pentylenetetrazol
Re: reuniens thalamic nucleus
RSA: retrosplenial agranular cortex
Rt: reticular thalamic nucleus
SC1: primary somatosensory cortex
SC2: secondary somatosensory cortex
sm: stria medullaris of the thalamus
SubI: subincertal nucleus
tFUS: transcranial focused ultrasound
TMS: transcranial magnetic stimulation
TX: ultrasound pulse transmit sequence
VA: ventral anterior thalamic nucleus
VC: visual cortex
vhc: ventral hippocampal commissure
VL: ventrolateral thalamic nucleus
VM: ventromedial thalamic nucleus
VNS: vagus nerve stimulation
ZI: zona incerta

## CRediT author statement

**Taewon Choi:** Investigation, Visualization, Writing—original draft, Writing—review & editing **Minseok Koo:** Investigation, Visualization, Writing—original draft, Writing—review & editing **Jaesoon Joo:** Methodology **Taekyung Kim:** Methodology **Youngmin Shon:** Methodology, Supervision, Writing—review & editing **Jinhyoung Park:** Conceptualization, Methodology, Visualization, Supervision, Writing—original draft, Writing—review & editing

## Acknowledgments

This work was supported in part by the National Research Foundation of Korea (NRF) grants funded by the Korea government (MIST) NRF-2020R1A2C2011808 (J.P.) and NRF-2021R1A4A1028713 (J.P.).

## Availability of data and materials

The authors declare that the data supporting the findings of this study are available within the paper and Supplementary Information files. Correspondence and requests for materials should be addressed to Jinhyoung Park and Young-Min Shon.

