## Supplementary Materials for "Rebound Excitation of Epileptiform Activities by Transcranial Focused Ultrasound Stimulation"

**This PDF file includes:**

Supplementary Figs. 1 to 2

Supplementary Table. 1

Supplementary Vids. 1 to 7

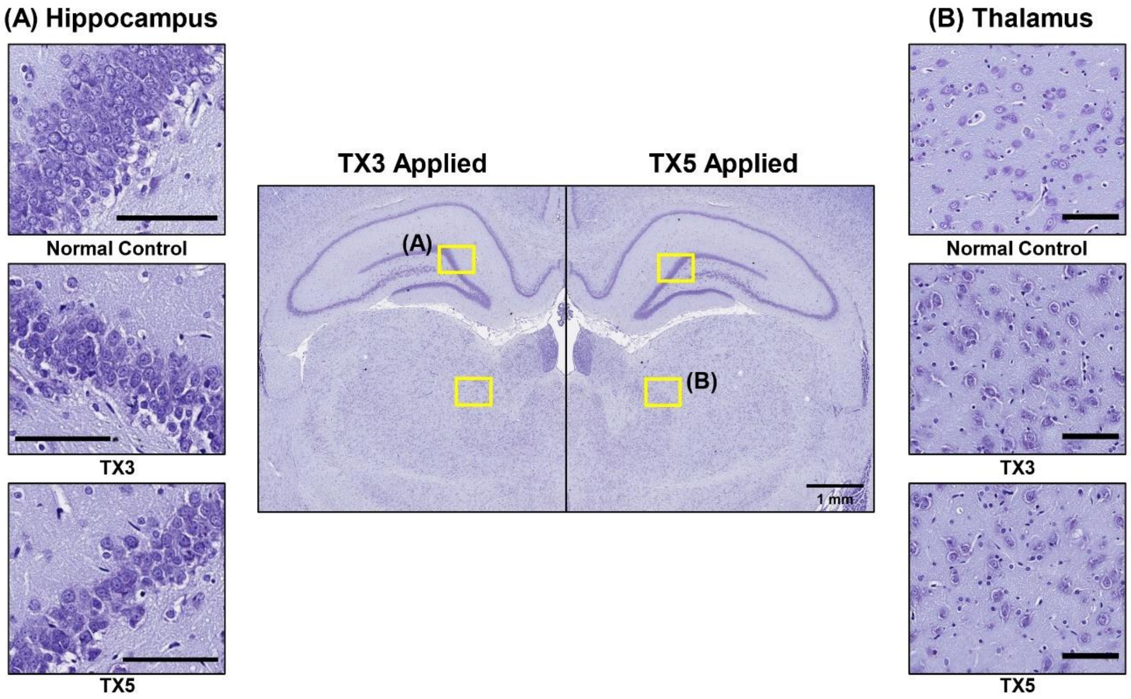


**Supplementary Fig. 1 Cresyl Violet staining analysis results.** (**A**) Representative images of Nissl-stained brain slice at the hippocampus. (**B**) Representative images of Nissl-stained brain slice images at the thalamic region. The Cresyl Violet Staining analysis demonstrated no damages in the rat brain after both TX3 and TX5 ultrasonic stimulation. The scale bar denotes 100 μm for all magnified images.

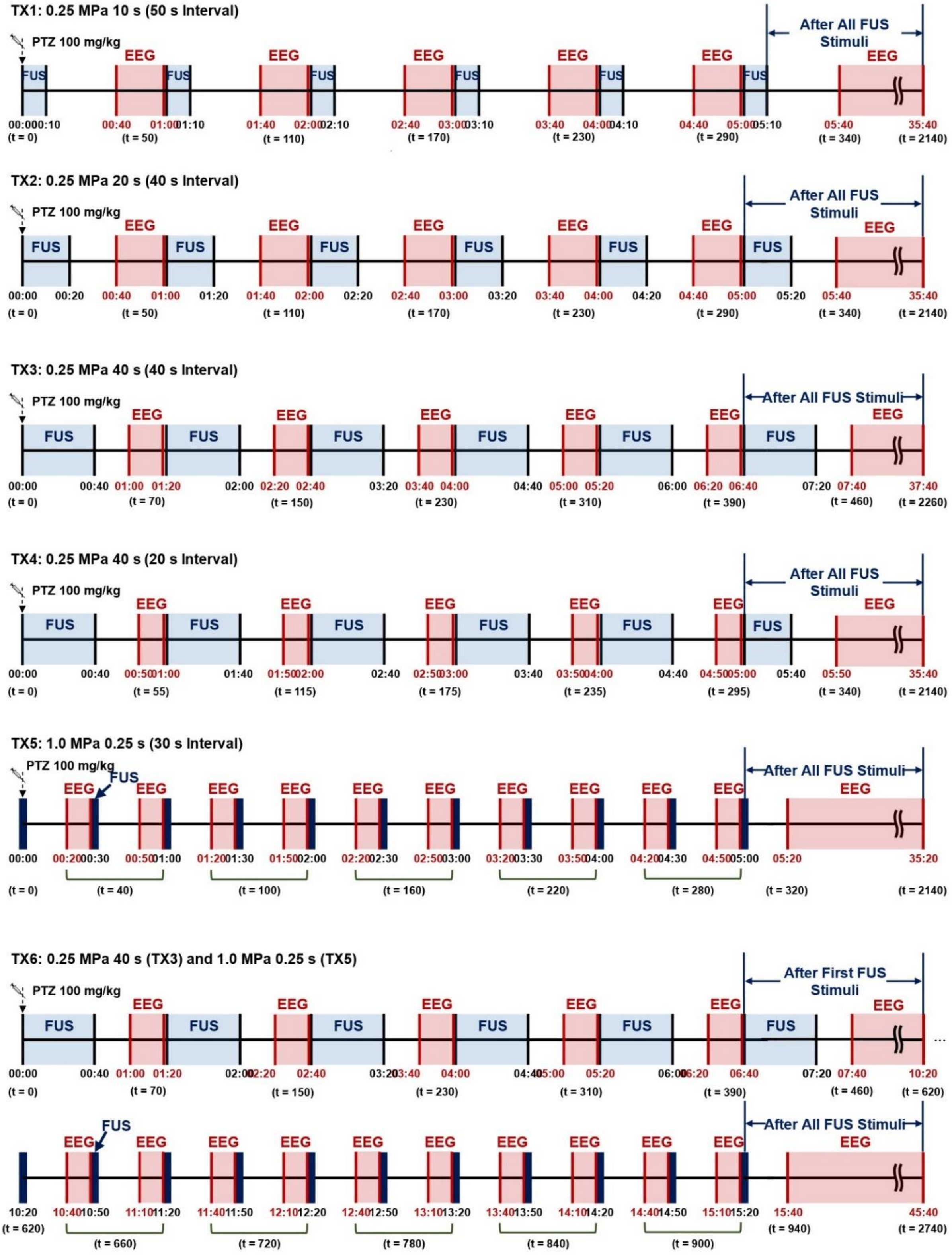


**Supplementary Fig. 2 Timeline of tFUS stimulation and EEG recording for all transmit condition groups.**

| Conditions | Acoustic Pressure | Single Stimulus Duration (SSD) | Stimulus Interval (Δ) | # Stim | Duty Cycle [%] | $I_{SPPA}$ [W/cm$^2$] | $I_{SPTA}$ [mW/cm$^2$] | Mechanical Index (MI) |
|---|---|---|---|---|---|---|---|---|
| **TX1** | 0.25 MPa | 10 s | Δ 50 s | 6 | 8.333 | 1.94 | 161.55 | 0.3 |
| **TX2** | 0.25 MPa | 20 s | Δ 40 s | 6 | 16.667 | 1.94 | 323.13 | 0.3 |
| **TX3** | 0.25 MPa | 40 s | Δ 40 s | 6 | 25 | 1.94 | 484.68 | 0.3 |
| **TX4** | 0.25 MPa | 40 s | Δ 20 s | 6 | 33.333 | 1.94 | 646.24 | 0.3 |
| **TX5** | 1.0 MPa | 0.25 s | Δ 30 s | 11 | 0.417 | 31.02 | 129.35 | 1.2 |
| **TX6** | 0.25 MPa / 1.0 MPa | 40 s / 0.25 s | Δ 40 s / Δ 30 s | 17 | 25 / 0.417 | 1.94 / 31.02 | 484.68 / 129.35 | 0.3 / 1.2 |

**Supplementary Table 1 Focused ultrasound parameters for each transmit condition, TX1-TX6.** The six transmit conditions are summarized in the table along with the calculated safety-related parameters. $I_{SPPA}$ denotes the spatial peak pulse average intensity, $I_{SPTA}$ denotes the spatial peak temporal average intensity, and #Stim denotes the number of stimulations for each transmit condition.

**Supplementary Vid. 1.**

2D topographical map representing the number of epileptic spikes ratio by time in PTZ-control.
https://drive.google.com/file/d/1ogPCFxQUUlMJf8HDiSdGhksbCtd_evh3/view?usp=sharing

**Supplementary Vid. 2.**

2D topographical map representing the number of epileptic spikes ratio by time in TX1.
https://drive.google.com/file/d/1zGYOjG_sF_UT-Fd4tFr1jGWTei5Mb3a7/view?usp=sharing

**Supplementary Vid. 3.**

2D topographical map representing the number of epileptic spikes ratio by time in TX2.
https://drive.google.com/file/d/10O71jtBsU9PDE0s0CFDX1cG4OywbQxbo/view?usp=sharing

**Supplementary Vid. 4.**

2D topographical map representing the number of epileptic spikes ratio by time in TX3.
https://drive.google.com/file/d/1qkkVAM8HqrcL_Hl-nIr7eQ6QLlGgHLWW/view?usp=sharing

**Supplementary Vid. 5.**

2D topographical map representing the number of epileptic spikes ratio by time in TX4.
https://drive.google.com/file/d/1O6JSum2M2ZbQ3VY0P6ZKGg25Kr71oHrV/view?usp=sharing

**Supplementary Vid. 6.**

2D topographical map representing the number of epileptic spikes ratio by time in TX5.
https://drive.google.com/file/d/1A31C8ZJeRNmraGdRHi116rnf0C1YLtju/view?usp=sharing

**Supplementary Vid. 7.**

2D topographical map representing the number of epileptic spikes ratio by time in TX6.
https://drive.google.com/file/d/1oZE9R6EI1fJAbSUBpFDGkhKGZnqDE5FU/view?usp=sharing
